# Kozak sequence acts as a negative regulator for *de novo* transcription initiation of newborn coding sequences in the plant genome

**DOI:** 10.1101/2020.11.28.402016

**Authors:** Takayuki Hata, Soichirou Satoh, Naoto Takada, Mitsuhiro Matsuo, Junichi Obokata

## Abstract

The manner in which newborn coding sequences and their transcriptional competency emerge during the process of gene evolution remains unclear. Here, we experimentally simulated eukaryotic gene origination processes by mimicking horizontal gene transfer events in the plant genome. We mapped the precise position of the transcription start sites (TSSs) of hundreds of newly introduced promoterless firefly luciferase (LUC) coding sequences in the genome of *Arabidopsis thaliana* cultured cells. The systematic characterization of the *LUC*-TSSs revealed that 80% of them occurred under the influence of endogenous promoters, while the remainder underwent *de novo* activation in the intergenic regions, starting from pyrimidine-purine dinucleotides. These *de novo* TSSs obeyed unexpected rules; they predominantly occurred ~100 bp upstream of the *LUC* inserts and did not overlap with Kozak-containing putative open reading frames (ORFs). These features were the output of the immediate responses to the sequence insertions, rather than a bias in the screening of the *LUC* gene function. Regarding the wild-type genic TSSs, they appeared to have evolved to lack any ORFs in their vicinities. Therefore, the repulsion by the *de novo* TSSs of Kozak-containing ORFs described above might be the first selection gate for the occurrence and evolution of TSSs in the plant genome. Based on these results, we characterized the *de novo* type of TSS identified in the plant genome and discuss its significance in genome evolution.

## INTRODUCTION

The process via which genetic novelty emerges has been a fundamental question of evolutionary biology. Because of the advancement of comparative genomics, our knowledge of new gene origination has been expanded; genes can be generated through the “bricolage” of pre-existing genetic materials, or can be originated *de novo* from non-coding DNA (Kaessmann, 2010; Cardoso-Moreira and Long, 2012; McLysaght and Guerzoni, 2015; Van Oss and Carvunis, 2019).

An essential question of gene birth is how newly originated gene sequences acquire their transcriptional competency, because it is a prerequisite for the mere sequences to become genes. Transcriptional competency is driven by a promoter, in which a specific sequence of elements and chromatin configuration exist for pre-initiation complex (PIC) binding and the initiation of transcription at a precise genomic position (Haberle and Stark, 2018; Andersson and Sandelin, 2020). As promoters activate the transcription of downstream DNA sequences, their evolution should be intrinsically connected to the functionalization of new genes. Comparative genomics has revealed that evolutionarily young genes acquired their transcriptional competency through (1) the utilization of duplicated ancestral promoters, (2) hijacking of pre-existing genes, promoter-like elements or spurious transcription units or (3) *de novo* emergence through mutations (Kaessmann, 2010; Li, Lenhard and Luscombe, 2018; Van Oss and Carvunis, 2019; Zhang *et al*., 2019). However, the promoters of such evolutionarily young genes are not so “young”, as they had been fixed in the genome through natural selection over a certain evolutionary period. Therefore, little knowledge is available regarding how newly originated coding sequences are transcribed and start evolving after their birth.

Experimental evolution is another approach to scrutinize such gene evolutionary processes, as it enables the analysis of “truly young” genes by mimicking the process of new gene origination in the native genomic environment (Garland, 2009). In plants, exogenously introduced coding sequences that mimic the originated genes through horizontal or endosymbiotic gene transfer (HGT/EGT) events have provided insights about how such newborn coding sequences acquire transcription ability. The escape of plastid DNA to the nucleus suggests that transferred plastid genes become transcriptionally active by trapping neighbouring eukaryotic promoters or by utilizing the prokaryotic plastid promoter sequences (Stegemann and Bock, 2006; Wang *et al*., 2014). By introducing promoterless coding sequences into the genome, promoter/gene-trapping screening also simulates gene origination processes (Friedrich and Soriano, 1991; Springer, 2000). A recent study reported that newly inserted promoterless coding sequences were transcribed without trapping any endogenous genes or transcription units, which indicated the origination of brand-new promoters in the plant genome (Kudo, Matsuo, and Satoh *et al*., 2020). However, the throughput of these studies was too limited to illustrate the general features of how newborn genes acquire transcriptional competency.

Here, we experimentally simulated gene origination processes in the plant genome to elucidate the manner in which newborn genes become transcriptionally active shortly after their birth. To overcome the low-throughput drawback of promoter/gene-trapping experiments, we previously applied a massively parallel reporter assay (Inoue and Ahituv, 2015) to the conventional promoter-trapping screenings, and established transgenic *Arabidopsis thaliana* T87 cell lines individually harbouring promoterless *LUC* open reading frames (ORFs) (Satoh and Hata *et al*., 2020). Based on the precise mapping of *LUC*-TSSs, we identified *de novo* TSSs; they occurred *de novo* about 100 bp upstream of the inserted coding sequences with specific avoidance of pre-existing putative ORFs containing a Kozak motif. We speculated that these features might reflect a first selection gate for the occurrence and evolution of *de novo* TSSs in the genome, regardless of the functionality of the newborn transcripts. Based on these results, we characterized the *de novo* TSSs detected in the plant genome and discuss their significance in genome evolution.

## RESULTS

### TSS determination for the newly inserted promoterless *LUC* genes

To scrutinize the manner in which newborn promoters occur in the plant genome, we analysed the transcription start sites (TSSs) of previously established transgenic *A. thaliana* T87 cell lines that harboured exogenously inserted promoterless luciferase (*LUC*) genes (Satoh and Hata *et al*., 2020). Because the cells experienced only 5–10 somatic cell divisions without luciferase-based screening, we assumed that they had retained the characteristic features of newborn genes.

To determine the TSSs and insertion loci of promoterless *LUC* genes in parallel, we modified the Cap-trapper method (Takahashi *et al*., 2012; Murata *et al*., 2014) for compatibility with inverse PCR for the selective analysis of the *LUC* transcripts. As shown in Figure 1a, we added the recognition sites of a rare-cutter enzyme at both ends of full-length cDNAs, to circularize them. *LUC* cDNAs were then selectively amplified by inverse PCR and subjected to paired-end deep sequencing. To obtain a precise map of *LUC*-TSSs and their corresponding insertion loci with single-nucleotide resolution, we carefully eliminated sequence artefacts derived from non-specifically amplified endogenous cDNAs and erroneous reads generated during the library preparation and sequencing steps (Figures S1 and S2, and Methods S1).

**Figure 1.**
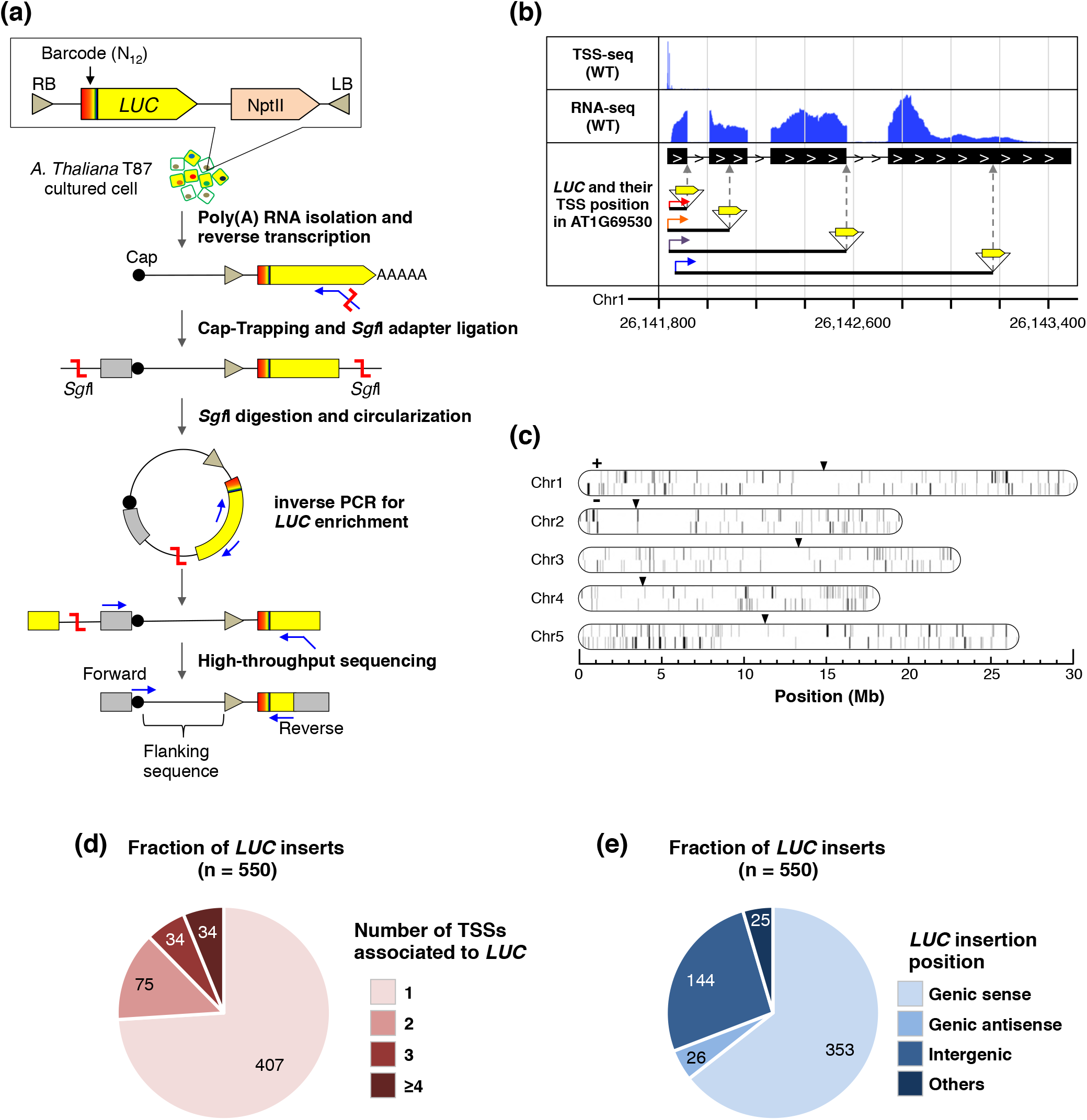
Determination of the TSSs of promoterless *LUC* genes at single-nucleotide resolution. (**a**) Experimental design of the parallel determination of promoterless *LUC* insertion sites and their corresponding TSSs. cDNAs reaching the 5’ end of *LUC* RNAs were prepared by the Cap-trapper method followed by inverse PCR. Amplified cDNAs were subjected to paired-end sequencing. For details, see the Methods. (**b**) Example of determined *LUC*-TSSs in the genome viewer. The coloured arrows indicate the determined *LUC*-TSSs. (**c**) Chromosomal map of all determined *LUC*-TSSs. The ticks indicate the genomic loci of 858 *LUC*-TSSs with sense (+) and antisense (−) orientations on *Arabidopsis thaliana* chromosomes. The black triangles indicate centromeres. (**d**) Relative abundance of the *LUC* inserts associated with the indicated number of TSSs. (**e**) Relative abundance of the *LUC* inserts with respect to the insertion types. Genic, protein-coding gene; Others, TAIR10-annotated region excluding protein-coding genes; Intergenic, unannotated region in TAIR10.

Figure 1b shows an example of the *LUC*-TSSs identified here, indicating that four independent *LUC* genes were inserted into the same gene body (AT1G69530), with their corresponding TSSs overlapping endogenous TSSs (Figure 1b). In total, we identified 550 *LUC* inserts and 858 corresponding TSSs across the *A. thaliana* genome (Figure 1c). Among the 550 *LUC* inserts, 74% were associated with a single TSS and the remainder were associated with two or more TSSs (Figure 1d). The *LUC* inserts were unbiasedly distributed over the *A. thaliana* genome (Satoh and Hata *et al*., 2020), whereas the *LUC* loci identified in this TSS analysis were over-represented in the genic regions (Figure 1e). This bias might reflect the fact that the inserts in the genic regions have relatively higher transcription levels and that their cDNAs were more easily obtained than were those located in intergenic regions. Nevertheless, we should note that one-fourth of the *LUC* inserts identified here were transcriptionally activated in the intergenic regions (Figure 1e) and were treated as candidate *de novo*-activated transcripts.

### *LUC*-TSSs were categorized into two types

To elucidate the mechanism via which promoterless *LUC* genes acquired their transcriptional competency, we next examined if the identified *LUC*-TSSs were associated with inherent TSSs. To prepare reference TSS datasets of WT cells, we performed genome-wide TSS-seq. We obtained 636,507 loci of highly reliable WT-TSS data, which covered 65.9% (18,064/27,416) of the annotated *A. thaliana* protein-coding genes. Compared with WT-TSSs, 64.6% (554/858) of the *LUC*-TSSs matched WT-TSSs with one-nucleotide resolution (Figure 2a). It was plausible to conclude that these *LUC*-TSSs were the result of transcriptional fusions with the endogenous transcripts. However, it was unclear whether the remaining *LUC*-TSSs were all *de novo* activated. To address this question, we tested the distribution of *LUC*-TSSs against the distance from the nearest WT-TSSs. Unexpectedly, the plot showed one clear inflection point at ±15 bp (Figure 2b). This result led us to hypothesize that a region of ±15 bp of WT-TSSs was under the influence of endogenous promoter activities. Based on these findings, we classified the *LUC*-TSSs into two categories; those located within ±15 bp of WT-TSSs and those located outside these regions. According to this categorization, out of 858 *LUC*-TSSs, we found that 654 (76%) were transcribed by pre-existing promoter activities, whereas the remainder (204, 24%) were candidate *de novo* TSSs that were unaffected by WT promoters (Figure 2c).

**Figure 2.**
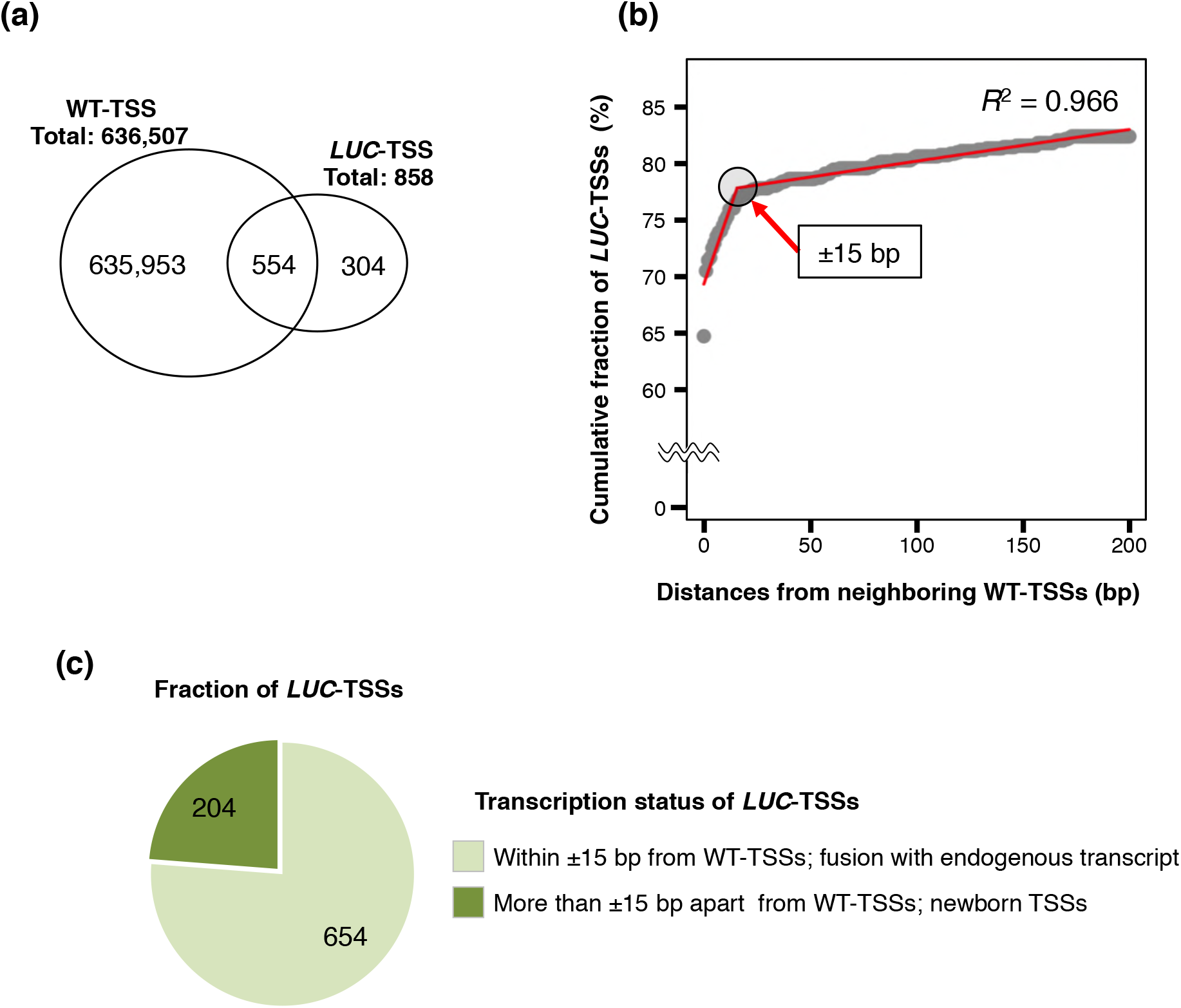
Categorization of *LUC*-TSSs with respect to WT-TSSs. (**a**) Venn diagram summarizing the overlap between the positions of WT-TSSs and *LUC*-TSSs at single-nucleotide resolution. (**b**) The grey dots show a cumulative fraction of *LUC*-TSSs according to the distances from their nearest WT-TSSs. The red line indicates the linear approximation of the grey dot plots, and the estimated inflection point is indicated by a black circle. The adjusted *R*^2^ was 0.966. (**c**) Number of *LUC*-TSSs categorized according to (b).

### Systematic classification of *LUC*-TSSs revealed the transcriptional activation mechanism of newborn genes

To clarify the features of *LUC*-TSSs in greater detail, we further classified them based on the combination of (i) *LUC* loci relative to the WT genes, (ii) TSS loci relative to the WT genes and (iii) types of *LUC*-TSS initiation (Figure 2c), to give 72 TSS types (Figure 3a). Among these 72 types, we identified 17 types in this study (Figure 3b, and Figure S3). This classification revealed that ~80% of the *LUC*-TSSs identified in this study were accounted for by transcriptional activation via the trapping of endogenous genes or transcription units (Figure 3b, and Figure S3). We found that transposable elements were also sources of transcriptional activation (Figure S3).

**Figure 3.**
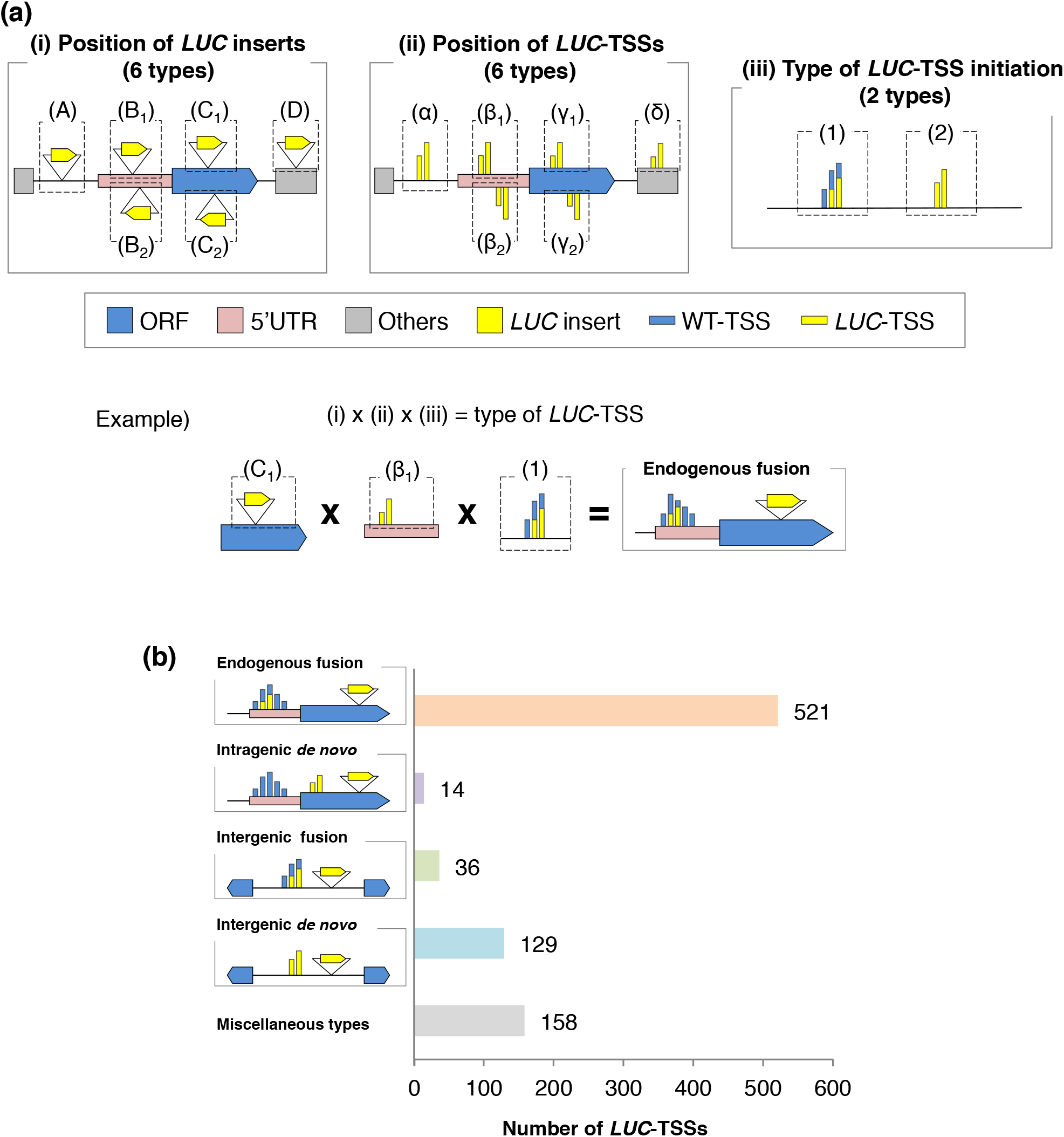
Classification of *LUC*-TSSs according to the combination of their genomic loci and types of TSS initiation. (**a**) *LUC*-TSSs were classified according to the combination of the position of (i) the *LUC* insert, (ii) the *LUC*-TSS relative to *Arabidopsis thaliana* annotated genes and (iii) the types of *LUC*-TSS initiation, as categorized in Figure 2c. Example showing the classification scheme of the “Endogenous fusion” type, in which the *LUC* gene was inserted in an endogenous ORF and the TSS initiated from the 5’-UTR of the ORF with an overlapping WT-TSS. (**b**) Number of *LUC*-TSSs of the representative insertion types, as described in the text. The contents of miscellaneous types are shown in Figure S3.

As our interest lay in the mechanism via which new promoters emerge in the plant genome, hereafter we focused on the *de novo*-activated TSSs in the intergenic regions (“Intergenic *de novo*”, A-α-2 type in Figure 3a). To compare the features of *de novo*-activated TSSs with those of pre-existing ones, we chose two additional types of *LUC*-TSSs: “Endogenous fusion” (C_1_-β_1_-1 type in Figure 3a), in which *LUC* genes were inserted in the pre-existing protein-coding genes and their TSSs overlapped with inherent WT-TSSs; and “Intergenic fusion” (A-α-1 type in Figure 3a), in which *LUC* genes were found in the intergenic region, but their TSSs overlapped with endogenous intergenic transcripts. In addition, we selected the “Intragenic *de novo*” type (C_1_-γ_1_-2 type in Figure 3a) to examine the differences in *de novo* TSSs between genic and intergenic regions. These four types accounted for 80% of the total *LUC*-TSSs identified here (Figure 3a).

### Newly activated TSSs have RNA polymerase II initiator and TATA-like motifs

Generally, transcription initiates preferentially at purine nucleotides (A/G) that are preceded by pyrimidine nucleotides (C/T) in the eukaryotic genome (Yamamoto *et al*., 2009; Haberle and Stark, 2018; Andersson and Sandelin, 2020). We confirmed that the *A. thaliana* protein-coding genes utilized the same initiation dinucleotide motif based on the TSS-seq of WT cells (Figure 4a, left and middle panels). We found that *LUC*-TSSs also initiated at a Py-Pu dinucleotide motif, even in the *de novo*-activated cases (Figure 4b–e, middle panels). A nucleotide composition analysis revealed the existence of an AT-rich region at ~30 bp upstream of *LUC*-TSSs, which might act as a TATA-box for facilitating PIC recruitment (Figure 4a–e, left panels). In addition to the AT-rich region described above, we were unable to find any characteristic motifs associated with the *de novo* TSSs.

**Figure 4.**
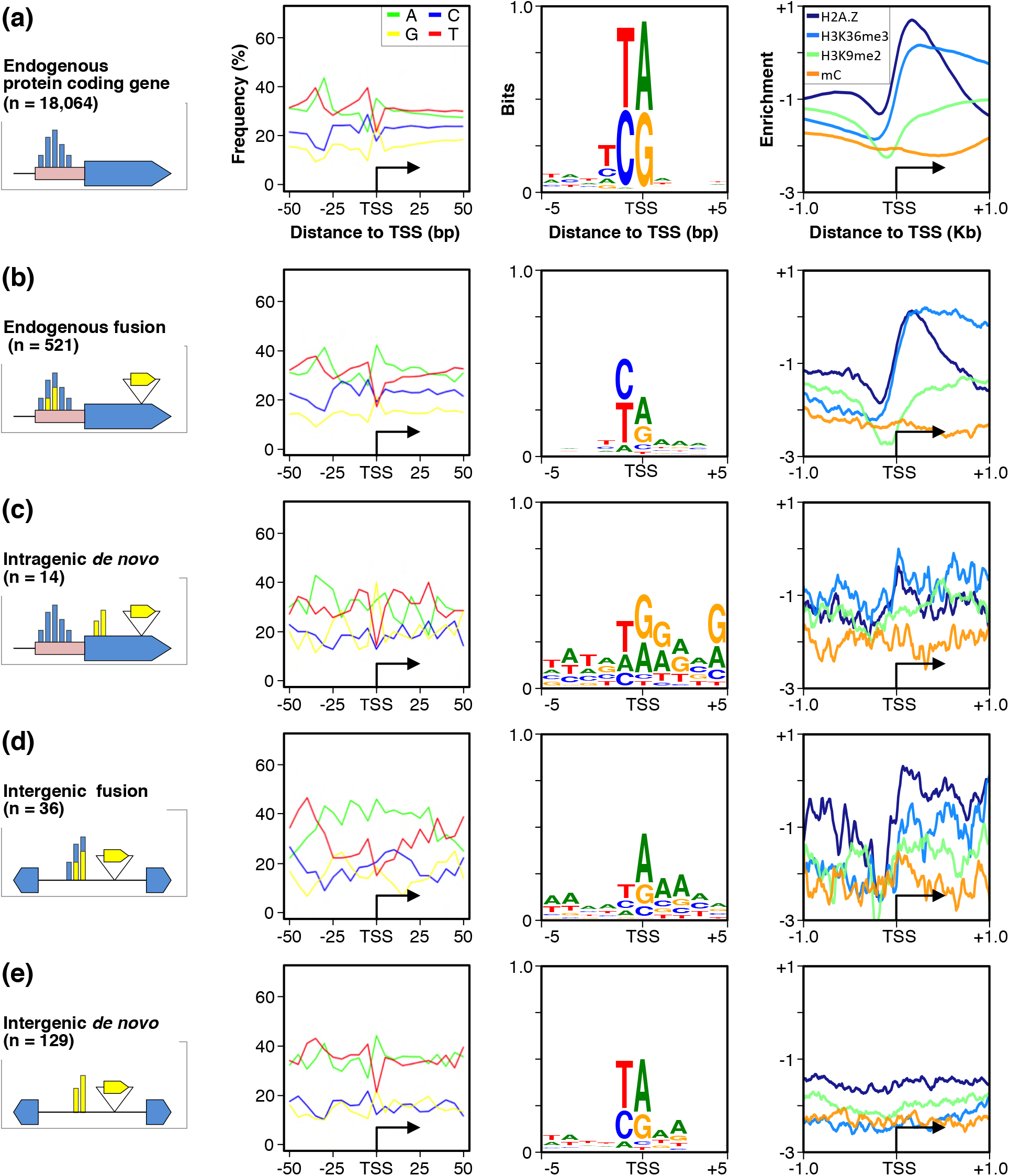
Sequence and epigenetic characteristics of the *LUC*-TSSs. (**Left panels**) Nucleotide frequency at 5 nt resolution centred on the TSSs of (**a**) endogenous protein-coding genes (n = 18,064) and *LUC*-TSSs classified as (**b**) “Endogenous fusion” type (n = 521), (**c**) “Intragenic *de novo*” type (n = 14), (**d**) “Intergenic fusion” type (n = 36) and (**e**) “Intergenic *de novo*” type (n = 129). The black arrows indicate the TSS. (**Middle panels**) Sequence logo around ±5 bp of the TSSs of (a) endogenous genes and (b–e) *LUC* genes. (**Right panels**) Distribution profiles of H2A.Z, H3K36me3, H3K9me2 and methylated cytosine (mC) in WT cells, within ±1.0 kb of the TSSs of (a) endogenous genes and (b–e) *LUC* genes.

### Promoter-like epigenetic status is not necessary for *de novo* TSS occurrence

Epigenetic status, including histone modification, histone variants, and DNA methylation, plays an important role in eukaryotic gene expression regulation (Gibney and Nolan, 2010). Therefore, we wondered whether the inherent epigenetic status is responsible for *LUC*-TSS activation. We first prepared a genome-wide map of four epigenetic marks in WT T87 cells, i.e., variant of histone H2A (H2A.Z) and lysine (K) tri-methylation of histone H3 (H3K36me3) as active transcription marks and lysine di-methylation of histone H3 (H3K9me2) and methylated cytosine (mC) as repressive marks, in the *A. thaliana* genome (Lauria and Rossi, 2011). In WT cells, we observed typical distributions of these four epigenetic marks around the TSS of endogenous protein-coding genes; H2A.Z exhibited peaks just downstream of TSSs, and H3K36me3, H3K9me2 and mC were distributed broadly along gene bodies (Figure 4a, right panel). The epigenetic landscapes of the “Endogenous fusion” type around its TSSs were similar to those of WT-TSSs (Figure 4a and b, right panels), because this type utilized the WT-TSS. In the “Intragenic *de novo*” type, slight enrichments of H2A.Z and H3K36me3 were found around the TSSs (Figure 4c, right panel). However, these apparent enrichments were attributed to those located upstream of WT-TSSs, because WT- and *LUC*-TSSs were located in the close proximity of this insertion type (Figure S4). We also found promoter-specific epigenetic patterns in the “Intergenic fusion” type, indicating that unannotated WT transcription was trapped in this case (Figure 4d, right panel). In contrast with these observations, no significant epigenetic patterns were detected around “Intergenic *de novo*” TSS loci (Figure 4e, right panel). Therefore, we concluded that a promoter-like epigenetic status was not necessary for the activation of *de novo* TSSs.

### *De novo* TSSs originated ~ 100 bp upstream of newborn coding sequences

Pervasive and spurious transcription is a characteristic of the eukaryotic genome and is one of the resources used for the transcriptional activities of new genes (Zhang *et al*., 2019). Our next question pertained to whether the *de novo* TSSs were activated by trapping cryptic transcripts that were not detected in our transcriptomics analysis of WT cells. To address this question, we attempted to determine the genomic distances between *LUC* insertion sites and the corresponding TSSs (TSS-to-*LUC* distances) for each TSS type. If the pre-existing WT-TSSs were utilized for *LUC*-TSSs after the insertion of *LUC* genes, the TSS-to-*LUC* distances should vary according to their insertion sites relative to the WT-TSSs. Expectedly, the TSS-to*-LUC* distances in these cases were broadly distributed (Figure 5a). Next, we examined the *de novo* TSSs. Surprisingly, “Intergenic *de novo*” TSSs initiated predominantly in the close vicinity of *LUC* insertion sites (median distance, 108 bp) (Figure 5a), with a relatively small coefficient of variation (CV = 0.60) compared with the “Intergenic fusion” type (CV = 1.08). This short and sharp distribution of TSS-to-*LUC* distances in the case of *de novo* TSSs was not explained by the size of the 5’ upstream intergenic regions of the inserts, because their sizes exhibited a large variation (Figure 5b, and Figure S5). We confirmed these distribution profiles in three different biological samples (Figure S6). Taken together, the unique features of *LUC*-to-*de novo* TSS distances suggest that they were not caused by the trapping of pre-existing cryptic transcripts at certain genomic loci; rather, the *de novo* TSSs were really caused by the *de novo* insertion of *LUC* coding sequences in their close proximity.

**Figure 5.**
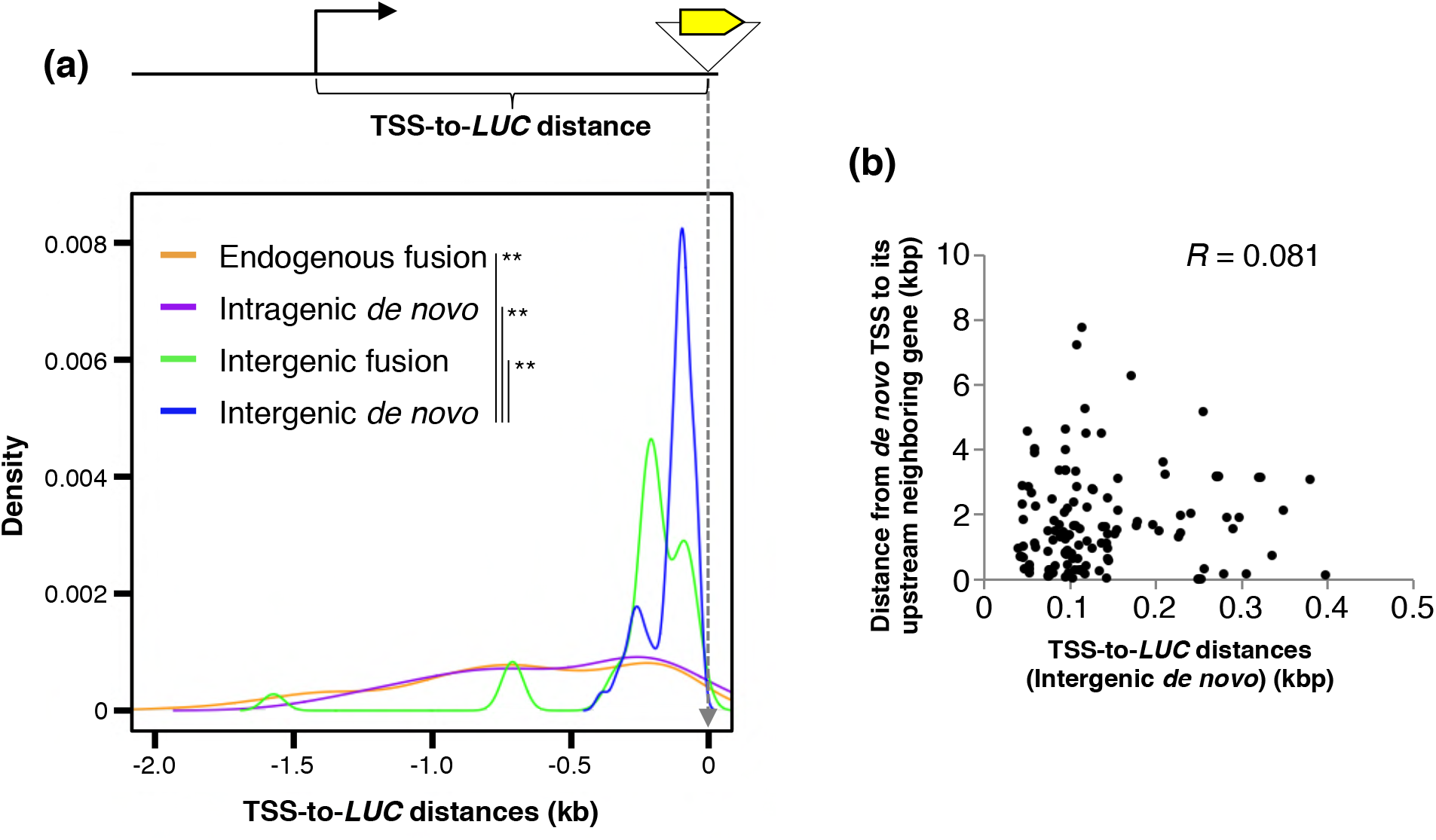
*De novo* TSSs occur in the 5’ proximity of the *LUC* inserts. (**a**) Density plot showing the distribution of the distance between the TSS and *LUC* insertion site (TSS-to-*LUC* distance) in each *LUC*-TSS type. The median TSS-to-*LUC* distance was (in bp): Endogenous fusion, 666; Intragenic *de novo*, 573; Intergenic fusion, 212.5; and Intergenic *de novo*, 108. \*\**P*-value < 0.01 (Wilcoxon rank-sum test). (**b**) Scatter plot showing the correlation between the TSS-to-*LUC* distance (“Intergenic *de novo*” type) and the TSS-to-upstream neighbouring gene distance. *R*, Pearson’s product-moment correlation test.

### *De novo* TSSs do not occur in the pre-existing Kozak-containing ORFs

In this study, *LUC* transcripts were translatable because they had a 5’-cap, a coding sequence and a 3’-polyadenylated tail. We wondered whether a relationship existed between this property and the *de novo* transcriptional activation. We observed that the initiation codon (ATG-triplets) frequency was low around *de novo* TSS loci compared with the distal regions (Figure 6a, and Figure S7). This characteristic was similar to the 5’-UTR of endogenous genes (Kim *et al*., 2007), which suggests that the *de novo* TSS regions might serve as the 5’-UTR of *LUC* messages. However, the determined *LUC* inserts did not have a minimum Kozak motif (A/GNN<u>AUG</u>G) (Nakagawa *et al*., 2008), as purine residue (A/G) was not enriched at the −3 position from the initiation codon of *LUC*-ORF (Figure 6b, and Figure S8a). In addition, the pre-existing putative ORFs around *de novo* TSS regions did not contribute to the translatability of the *LUC* messages; such putative ORFs provided an in-frame Kozak-ATG to the downstream *LUC*-ORFs in only 6.9% of cases (9/129) (Figure S8b). These results indicate that our *LUC*-TSS population was not enriched for translatability of the *LUC* messages. This was a reasonable conclusion because transgenic cells had not been screened for luciferase activity. However, we found that Kozak-containing ORFs exhibited an unusual distribution around *de novo* TSSs: these two entities were mutually exclusive (Figure 6c and d). As shown in Figure 6c, *de novo* TSSs did not occur within Kozak-containing ORFs (Figure 6c, middle panel, and Figure S8c), while ORFs without Kozak sequences were uniformly distributed around *de novo* TSS loci as well as in randomly sampled intergenic regions (Figure 6d, left and middle panels). These distribution patterns were commonly observed among three distinct biological replicates (Figure S8d). Interestingly, the repulsion between TSSs and ORFs was more evident in WT genes, with few ORFs found around TSSs and 5’-UTRs regardless of the Kozak motif (Figure 6c and d, right panels). Therefore, the anti-Kozak rule of the *de novo* TSSs might be an initial stage of the repulsion between the TSSs and ORFs. These findings imply that the anti-Kozak rule might be an outcome of the immediate responses to sequence insertion, with subsequent natural selection steps eliminating the ATG-triplets interposed in the 5’-UTR through evolutionary timescales.

**Figure 6.**
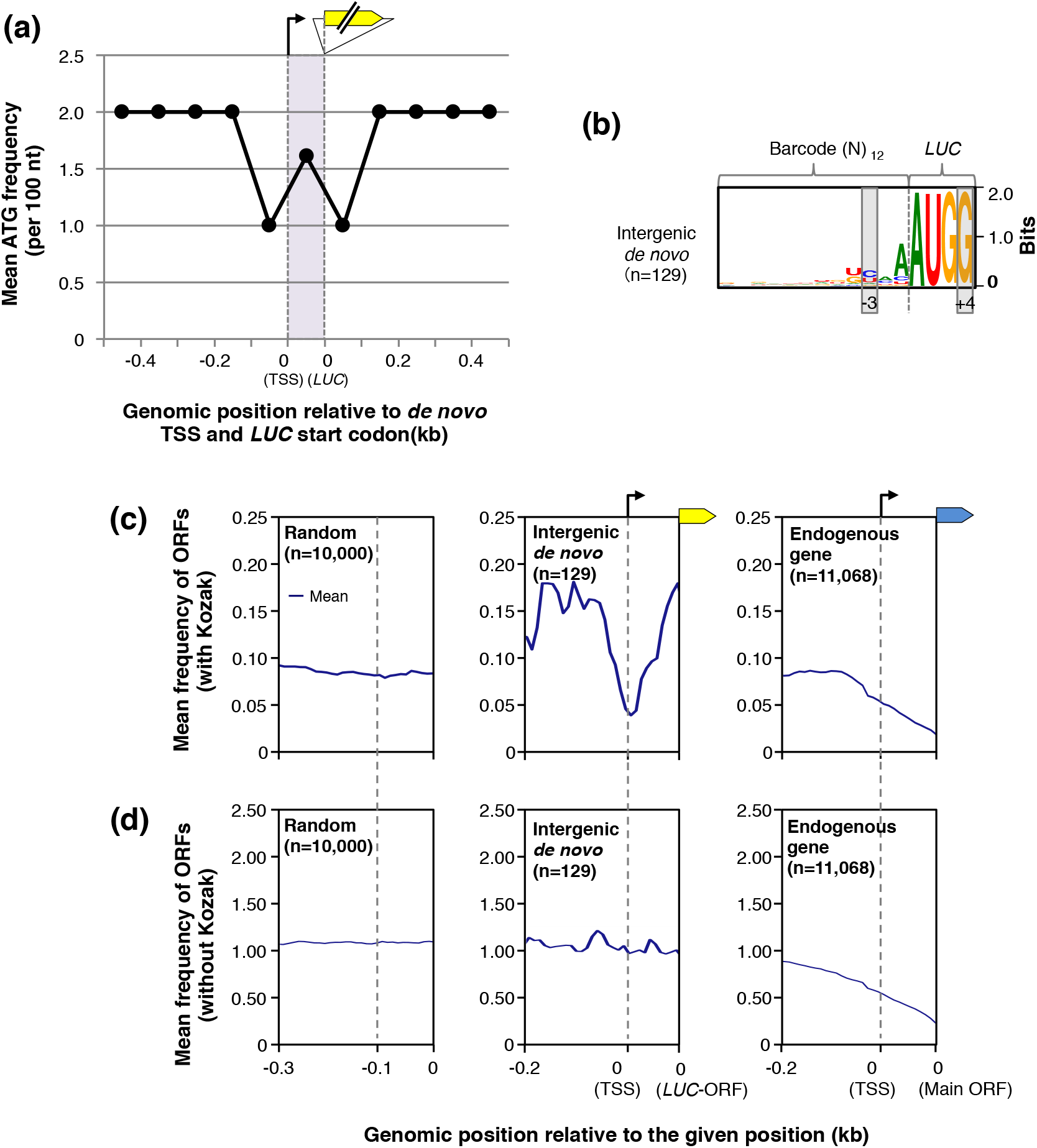
*De novo* TSSs avoid pre-existing Kozak-containing ORFs. (**a**) Mean frequency of the initiation codon (ATG) per 100 bp around *de novo* TSS regions. The ATG frequency in the *de novo* TSS regions was normalized per 100 bp. (**b**) Sequence logo of the barcode region on the Intergenic *de novo*-type *LUC* inserts (n = 129). The conserved positions of a minimum Kozak motif (A/GNNAUGG) are indicated by the grey boxes. (**c** and **d**) Meta-plot of the distribution profiles of ORFs (**c**) with or (**d**) without a Kozak motif within 0.3 kb of randomly sampled intergenic regions (left panels), the region from 0.2 kb upstream of the Intergenic *de novo* TSS to its *LUC*-ORF (middle panels) and the region from 0.2 kb upstream of the TSS of endogenous protein-coding genes to their main ORF (right panels). The frequencies of ORFs located within the region from the *de novo* TSS to the *LUC*-ORF and from the genic TSS to the main ORF were normalized per 0.1 kb. *Arabidopsis thaliana* genes with introns in the 5’-UTR were excluded from the analysis. The grey dotted lines indicate the TSS positions.

## DISCUSSION

A long-standing question in biology concerns the principles of evolutionary innovation. The origination of new genes is a central driver of evolution and has attracted the interest of researchers. Comparative genomics has been an effective tool in this research area, as it has provided various insights into the gene evolutionary process (Kaessmann, 2010; Cardoso-Moreira and Long, 2012; McLysaght and Guerzoni, 2015; Van Oss and Carvunis, 2019). However, the time resolution of comparative genomics has intrinsic limitations and is not suitable for dissecting the ordered events of the gene origination process in a relatively short period. In this regard, our artificial evolutionary experiment, which mimicked the HGT/EGT process, has advantages in the study of a much nearer time point to gene birth. By attempting to perform an elaborate classification of the gene insertion types relative to the annotated gene loci (Figure 3, and Figure S3), we succeeded in isolating the genuine *de novo*-type transcription of the inserts and in discriminating it from the other types that occurred under the influence of pre-existing promoters.

*De novo* transcription had the following characteristics: (1) its TSS was located at a Py-Pu dinucleotide located ~100 bp upstream of the *LUC* insert; (2) it tended to have an AT-rich region located ~30 bp upstream of the TSS; (3) inherent promoter-like epigenetic profiles were not needed; and (4) its TSS avoided overlap with pre-existing Kozak-containing ORFs. These analyses were performed using transgenic cells that experienced only 5–10 vegetative cell divisions, and were not screened for luciferase activity (Satoh and Hata *et al*., 2020). Therefore, these characteristics were intrinsic properties of noticeably young promoters that were observed right after their birth, before their exposure to evolutionary selective pressures.

Based on the sequence characteristics of *de novo* TSSs mentioned above, as well as the 5’-capped and 3’-polyadenylated nature of the RNA samples (Figure 1a), it is probable that the *de novo* transcription that we detected in this study was mediated by RNA polymerase II (pol II) (Haberle and Stark, 2018; Andersson and Sandelin, 2020). An AT-rich region was not always detected upstream of the *de novo* TSS (Figure 4); hence, it does not seem to be necessary for *de novo* transcription, but likely facilitates chromatin opening (Zuo and Li, 2011). The relatively low GC content of the *A. thaliana* genome (36%) (Barakat, Matassi and Bernardi, 1998) might increase the occurrence of *de novo* TSSs.

As *de novo* TSSs occur without inherent promoter-like epigenetic profiles (Figure 4e), a transcription-supporting chromatin configuration in these cases is supposed to be formed after sequence insertion. We found analogous cases in transgenic plants, in which promoterless *LUC* genes became transcriptionally activated concomitant with chromatin remodelling around the *LUC* insertion loci (Kudo, Matsuo, and Satoh *et al*., 2020; Hata and Takada *et al*., 2020). From the massive analysis of transgenic cultured cells, we also found that transcriptional activation occurred stochastically at 30% of the insertion events across the genome and was independent of chromosomal loci, suggesting that this transcriptional activation reflects the stochastic nature of chromatin remodelling (Satoh and Hata *et al*., 2020). Taken together, these findings suggest that gene insertion events stochastically activate local chromatin remodelling to form a transcription-competent chromatin configuration. If this is the case, how is the inserted *LUC* ORF sequence involved in this phenomenon?

*De novo* TSSs occurred ~100 bp upstream of *LUC* ORFs (Figure 5a), suggesting that *LUC* ORFs are involved in the positioning of the PIC. This putative positioning mechanism is buttressed by our previous observation. When core promoter regions were triplicated in front of the *LUC* ORF, the most proximal core promoter unit was predominantly utilized in transgenic plants (Kudo, Matsuo, and Satoh *et al*., 2020). Therefore, the coding sequence is likely to act as a *cis*-determinant element of the pol II PIC recruitment. The mechanism underlying this PIC positioning warrants further analysis.

Another intriguing finding of this study was the mutual repulsion between the *de novo* TSSs and Kozak-containing ORFs (Figure 6c). The simplest explanation for this repulsion is that Kozak-containing ORFs are covered by transcription-repressive chromatin marks, as is known for many annotated genes (Neri *et al*., 2017; Nielsen *et al*., 2019). Notably, this repressive effect was not observed for ORFs without a Kozak motif (Figure 6d). Considering that the Kozak motif is generally thought to function on mRNA molecules, the repulsion detected here suggests that the epigenetic configuration of the genomic ORF is retro-regulated by the mRNA translatability. Does this feedback mechanism operate within the nucleus, or is it linked to cytoplasmic activities, as are the mRNA surveillance mechanisms (Chang, Imam and Wilkinson, 2007; Smith and Baker, 2015)? This question deserves further investigation.

Based on the collective findings reported above, we propose a model to explain the gene origination process in the plant genome (Figure 7). First, when brand-new coding sequences are originated/introduced by genome shuffling or the EGT/HGT process, initial transcriptional activation occurs stochastically anywhere in the genome (Figure 1c) (Satoh and Hata *et al*., 2020). The newly occurred TSSs avoid pre-existing Kozak-containing ORFs to avoid interference with the pre-existing genetic information (Figure 6c). These processes within a biochemical timescale determine the initial configuration of the pol II promoters, which are subjected to subsequent natural selection on genetic and evolutionary timescales. In this model, the initial recruitment steps of the transcriptional machinery warrant further investigation (Step 2 in Figure 7).

**Figure 7.**
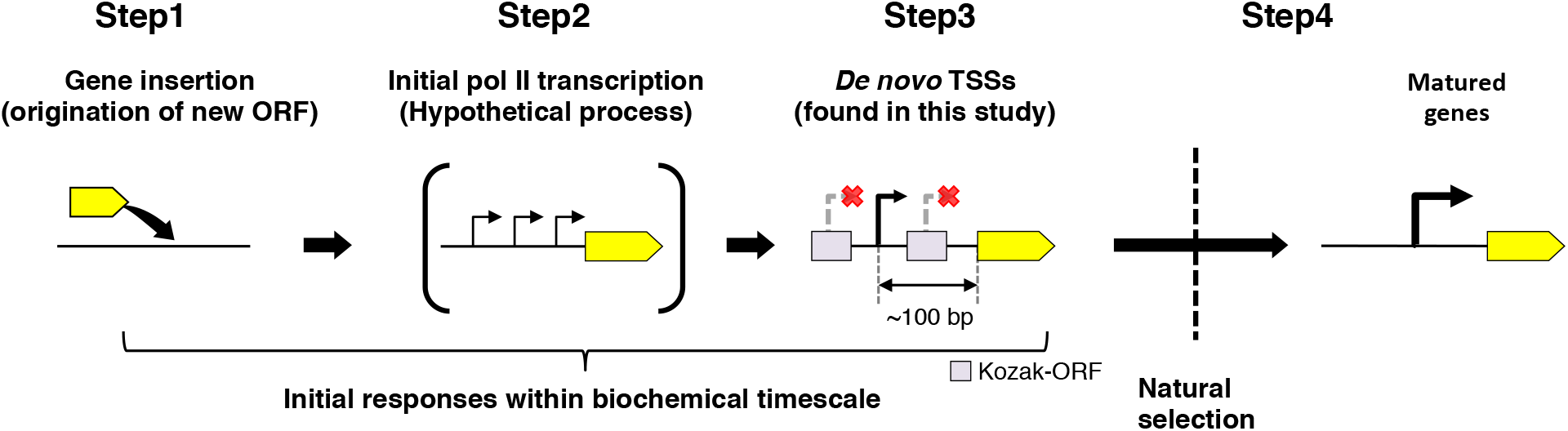
Model of the evolutionary processes of new genes. Brand-new coding sequences are originated/introduced by genome shuffling or the EGT/HGT process. *De novo* TSSs occur in response to the origination of a new coding sequence, with satisfying an anti-Kozak rule. *De novo* TSSs are originated within biochemical timescale, independently of the functionality of the messages. After *de novo* TSS occurrence, the neighbouring putative ORFs are eliminated via function-based natural selection in the evolutionary timescale.

In conclusion, our artificial evolutionary experiment allowed the detailed scrutiny of the origination process of functional genes in a biochemical timescale. We describe unique properties of *de novo* TSSs for the first time, which served as the basis of a gene origination model in the plant genome. Because the current study was performed using cultured cells, the genetic behaviour of *de novo* transcription requires further examination regarding heredity and functional adaptation with/without selective pressures.

## EXPERIMENTAL PROCEDURES

### Plant material and growth condition

*Arabidopsis thaliana* T87 cultured cells (Axelos *et al*., 1992) were maintained in mJPL3 medium (Ogawa *et al*., 2008) at 22°C with shaking under continuous-light conditions (50–70 μE m^−2^ s^−1^). One-week-old cultures were harvested using a 10 μm nylon mesh, washed with H_2_O twice and subjected to DNA, RNA and chromatin isolation, respectively. We set up two biological replicates for all further experiments, which were processed independently in each experiment.

### T87 wild-type (WT) TSS-seq library preparation

All primers used in this study are listed in Table S1. Total RNA was isolated from WT T87 cells using an RNeasy Plant Mini Kit (QIAGEN) followed by DNase I treatment. Next, polyadenylated RNA (poly (A) RNA) was enriched using a Dynabeads mRNA Purification Kit (Invitrogen) according to the manufacturer’s protocols. Poly (A) RNA (2.0 μg) was reverse transcribed using 1000 pmol of random hexamer primers tailed with an Illumina Rd1 adapter. Cap-trapping and subsequent adapter ligation (Illumina Rd2 adapter) steps were performed according to the published methods (Takahashi *et al*., 2012; Murata *et al*., 2014). Double-stranded cap-trapped cDNAs were amplified using a Nextera XT index primer (Illumina), then size selected at 200–400 bp using AMPure beads (BeckmanCoulter). Next-generation sequencing (NGS) was performed on an Illumina Mi-Seq platform using a 76 bp paired-end protocol.

### T87 WT TSS-seq data processing

Low-quality reads (Q30 <80%) were discarded using FASTX_Toolkit (http://hannonlab.cshl.edu/fastx_toolkit/). The first nucleotide of the forward reads was added by the library preparation step, and the second nucleotide was attributed to non-templated addition by reverse transcriptase. Therefore, these two nucleotides were trimmed from both ends and were used for TSS validation after mapping according to Yamamoto *et al*. (Yamamoto *et al*., 2009). Processed paired reads were mapped to the TAIR10 release of the *A. thaliana* genome assembly (https://www.arabidopsis.org/) using STAR (version: 2.5.4b) (Dobin *et al*., 2013) with the following parameters: *STAR −outFilterMultimapNmax 1 −alignEndsType EndToEnd – alignIntronMax 6000* (Marquez *et al*., 2012) –*twopassMode Basic*. Concordantly and uniquely mapped forward reads were extracted according to their SAM Flags (Li *et al*., 2009); 99 (sense to reference) and 83 (antisense to reference). Precise TSSs were called according to their cap signature (Yamamoto *et al*., 2009).

### T87 WT chromatin immunoprecipitation sequencing (ChIP-seq) library preparation

Chromatin isolation and subsequent ChIP of WT T87 cells were performed according to the published method (Satoh and Hata *et al*., 2020) with modifications, as follows. Fixed cells (0.2 g) were used for chromatin isolation. ChIP was performed with 10–20 ng of solubilized chromatin, Dynabeads Protein-G magnetic beads (Invitrogen) and antibodies: 2.4 μg of an anti-H2A.Z rabbit polyclonal antibody (Kudo, Matsuo, and Satoh *et al*., 2020) and 1.0 μg of an anti-H3K36me3 rabbit polyclonal antibody (Abcam: ab9050) were used in this experiment. Successful enrichment of ChIPed DNA was validated by quantitative PCR (qPCR) according to Deal *et al*. (Deal *et al*., 2007) for H2A.Z, and to Yang *et al*. (Yang, Howard and Dean, 2014) for H3K36me3. ChIP-seq libraries were prepared using a DNA SMART ChIP-seq Kit (Clontech) with 1.0 ng of ChIPed DNA and input DNA (DNA extracted from sheared chromatin), respectively. Libraries were size selected at 200–400 bp using AMPure beads. NGS was performed using a 51 bp single-ended protocol on an Illumina HiSeq 2000 platform.

### T87 WT methyl-CpG binding domain protein-enriched genome sequencing (MBD-seq) library preparation

DNA was extracted from WT T87 cells using a DNeasy Plant Mini Kit (QIAGEN). DNA (2.0 μg) was sheared to obtain 50–500 bp fragments (median size, 200 bp) by sonication (TOMY, UD-201), and purified using a QIAquick PCR Purification Kit (QIAGEN). Sheared DNA (500 ng) was used for methylated DNA enrichment, followed by NGS library preparation using an EpiXplore Meth-Seq DNA Enrichment Kit (Clontech). Methylated DNA enrichment was verified by qPCR according to Erdmann *et al*. (Erdmann *et al*., 2014). Enriched DNA (5.0 ng) was used for NGS library preparation. Libraries were size selected at 200–400 bp using AMPure beads. Sequencing was performed using a 51 bp single-ended protocol on an Illumina HiSeq 2000 platform.

### T87 WT ChIP-seq and MBD-seq data processing

ChIP-seq data for H3K9me2 were retrieved from *DDBJ Sequence Read Archive* under accession DRA009315. Low-quality reads (Q20 <80%) were discarded using FASTX_Toolkit (http://hannonlab.cshl.edu/fastx_toolkit/). The first three nucleotides added during the library preparation step were trimmed. Processed reads were mapped to the *A. thaliana* genome (TAIR10) using Bowtie2 (version: 2.2.5) (Langmead and Salzberg, 2012) allowing for one mismatch. Uniquely mapped reads were adopted, and duplicated reads were removed using Picard tools (version: 2.16.0) (http://broadinstitute.github.io.picard/).

### *LUC*-TSS-seq library preparation

Transgenic T87 cells harbouring promoterless *LUC* genes were established previously (Satoh and Hata *et al*., 2020). For three biological replicates of transformed cells, we prepared two technical replicates, respectively. RNA preparation, Cap-trapping and subsequent adapter ligation were performed as described for the WT TSS-seq library preparation with modifications, as follows (Figure 1a). Poly (A) RNA (2.0 μg) was reverse transcribed using a 0.2 μM *LUC*-specific primer tailed with an SgfI site. After Cap-trapping, the adapter oligo containing the SgfI site was ligated to the 3’ end of the cDNA. Subsequently, double-stranded cDNA (1–5 ng) was completely digested by SgfI. Because SgfI sites appear at an exceptionally low frequency in the *A. thaliana* genome (~2 sites/Mb), we could avoid undesirable digestion at endogenous SgfI sites almost completely. Digested cDNAs were then circularized by T4 DNA ligase, and 0.5–1 ng of circularized cDNA was used for inverse PCR to enrich *LUC* cDNA using a *LUC*-specific primer set. Subsequently, a sequencing library was prepared by two rounds of PCR; the first round was performed to add Illumina adapters, and the second was carried out using Nextera XT index primers. Libraries were sequenced on an Illumina MiSeq platform.

### *LUC*-TSS-seq data processing

Forward and reverse reads (TSS side and *LUC* side, respectively) were independently processed before mapping for the sake of removing cloning artefacts, trimming unmappable sequences derived from library design, and determining precise TSSs and their barcode sequences (Methods S1 and Figure S1). Subsequently, processed paired reads were mapped onto the *A. thaliana* genome (TAIR10) using STAR (version: 2.5.4b) (Dobin *et al*., 2013) with the following parameters: *STAR --outFilterMultimapNmax 1 –alignEndsType EndToEnd – alignIntronMax 6000* (Marquez *et al*., 2012) *–outFilterMismatchNoverLmax 0.06 twopassMode Basic*. Concordantly and uniquely mapped read pairs were collected according to their SAM Flag pairs (Li *et al*., 2009); the forward and reverse read sets were 99 and 147, or 83 and 163, respectively. Precise TSSs were called according to their cap signature (Yamamoto *et al*., 2009). Subsequently, we eliminated *LUC*-TSS artefacts caused by PCR and sequencing errors using the procedures described in Methods S1 and Figure S2.

### *LUC*-TSS classification

The distances between individual *LUC*-TSSs and their nearest WT-TSS in the same strand were calculated using bedtools (version: v2.17.0) (Quinlan and Hall, 2010). Using the distribution curve of *LUC*-TSSs against the distance described above, 1,000 times bootstrap repetition of linear approximation using the “segmented” R package (https://CRAN.R-project.org/package=segmented) revealed the presence of an inflection point at ±15 bp from the nearest WT-TSS. According to the inflection point, *LUC*-TSSs were divided into two groups: within or outside of ±15 bp from the nearest WT-TSS. *LUC*-TSSs were then classified according to the combination of TSS and *LUC* positions while considering their orientations (sense or antisense) relative to the *A. thaliana* genome annotations, as well as the initiation type of the *LUC*-TSSs grouped as described above. For genome annotation, we used the TAIR10 annotation with the exception of the 5’-untranslated region (5’-UTR); these regions were expanded to 200 bp upstream of the annotated position. The annotated regions, with the exception of protein-coding genes (i.e., transposable elements), were defined as “Others”.

### TSS characterization

Nucleotide frequency was calculated in a 5 bp window around ±50 bp of *LUC*-TSSs and WT-TSSs, respectively. The sequence logo was generated by the “RWebLogo” R package (version: 1.0.3) (https://CRAN.R-project.org/package=RWebLogo). A metagene plot of epigenetic status was generated by deeptools (version: 3.2.1) (Ramírez *et al*., 2014) using TAIR10 annotation and *LUC*-TSS positions, respectively. A motif enrichment analysis was performed using Centrimo with reported motif databases (Bailey and Machanick, 2012; O’Malley *et al*., 2016). Initiation codon (ATG) frequency was calculated in a 100 bp window around *de novo* TSSs and *LUC*-ORFs. The real lengths of the regions located between individual *de novo* TSSs and *LUC*-ORFs varied according to individual sites. Therefore, their individual lengths were normalized to 100 bp when calculating ATG frequency. The distribution of putative ORFs was analysed around ±0.2 kb of intergenic *de novo* TSSs, 5’-UTR of endogenous genes and randomly extracted intergenic regions, respectively. The 5’-UTR of endogenous protein-coding genes was defined as the region located between the annotated initiation codon and their strongest TSS, as determined by the TSS-seq analysis of WT cells. 5’-UTRs with splice sites were excluded from the analysis. Randomly extracted intergenic regions were prepared via the random extraction of 100 bp fragments from the intergenic region over 10,000 times. The heat map and meta-plot of ORF distribution were generated by deeptools (version: 3.2.1) (Ramírez *et al*., 2014).

## ACCESSION NUMBERS

Next generation sequencing data of WT T87 cells are available in the *DDBJ Sequenced Read Archive* under the accession number: DRA009316

## ACKNOWLEDGEMENTS

This work was supported by Japan Society for the Promotion of Science (JSPS) KAKENHI Grant Number: 26660008, 17J04887.

## SUPPORTING INFORMATION

Additional supporting information is found in the online version of this article.

**Figure S1.** Schematic illustration of paired-end read processing before mapping.

**Figure S2.** Mapping depth-based determination of genuine *LUC*-TSSs.

**Figure S3.** All determined *LUC*-TSS types.

**Figure S4.** Distribution of epigenetic marks around the TSS of Intragenic *de novo* type.

**Figure S5.** Comparison between the length of *de novo* TSS region and that of the corresponding intergenic region.

**Figure S6.** Distribution profiles of TSS-to-*LUC* distances in biological replicates.

**Figure S7.** Initiation codon frequency around *de novo* TSS region.

**Figure S8.** Distribution profiles of Kozak sequence around *LUC* insertion loci.

**Table S1.** Information of All DNA oligos used in this study.

**Methods S1.** Detailed methods for *LUC*-TSS data processing.

## Methods S1

### Forward read (TSS side) processing before mapping (Figure S1)

1. Read trimming Sequences were trimmed to 75 nt from the 3’ end of individual reads in order to remove low-quality sequences.
2. Cap identification and trimming The first two nucleotides were trimmed since they are added in the library preparation step and therefore unmappable to the genome. Note that the second nucleotide was corresponding to the 5’ cap position, so their sequence information was used for TSS validation as cap-signature (Yamamoto *et al*., 2009).
3. Check for overlapping sequences to reverse reads If the genomic position of *LUC*-TSS is very close to the *LUC* insert, forward read sequences would overlap their corresponding reverse read. For extracting properly mappable genomic sequences, we detected such genome-*LUC* chimeric junctions in individual forward reads by BLASTn (version: 2.4.0+) (Camacho *et al*., 2009) in megablast task (Morgulis *et al*., 2008) using *LUC* insert sequences obtained from the corresponding reverse read as a subject. Aligned sequences in forward reads were trimmed when they fulfill following cases; (1) the alignment started 5’ end of subject (from reverse read), (2) the alignment reached 3’ end of the query (forward read), (3) the alignment has no gaps, and (4) the alignment allowed up to 3 mismatches. If the above conditions were not satisfied, read trimming was not performed.
4. Extracting flanking sequences to map After checking and removing overlapping sequences to reverse reads, flanking genomic sequences were extracted for paired-end mapping.

### Reverse read (*LUC* side) processing before mapping (Figure S1)

1. Read trimming Sequences were trimmed to 100 nt from the 3’ end of the read in order to remove low-quality sequences. If the reverse read was shorter than 100 nt, the last two nucleotides were trimmed since they are derived from sequence library preparation steps.
2. Identification of genome-*LUC* chimeric junction In order to identify precise junction point between the genome and *LUC* insert, and also to identify each barcode sequence, each reverse read was aligned with ideal *LUC* insert sequence (5’-TTATGTTTTTGGCGTCTTCCATNNNNNNNNNNNNCTGTAAGCTGATAACGTCGAGGCCTTGA-3’; N corresponds to barcode) as a BLASTn (version: 2.4.0+) (Morgulis *et al*., 2008) subject sequence. Aligned sequences in reverse reads were trimmed when they fulfill following cases; (1) the alignment started 5’ end of both subject (ideal sequence) and query (reverse read), (2) the alignment length was at least 45 nt, (3) the alignment has no gaps, and (4) the alignment allowed up to 15 mismatches (this means 3 mismatches allowed excluding barcode sequences). If the above conditions were not satisfied, those reads (and also corresponding forward reads) were discarded as contaminated artifacts.
3. Extracting flanking sequences and barcode sequences After checking and removing *LUC* insert sequences, flanking sequences were extracted for paired-end mapping. In addition, each barcode sequence was extracted for *LUC* insert validation.

### Genuine TSS-to-*LUC* tag calling (Figure S2)

In order to eliminate aberrant *LUC*-TSS candidates caused by the mutations that occurred during PCR and sequencing steps, we made four histograms (see below) of mapping depth. Each mapping depth was calculated according to the read-tags specified by the TSS position, *LUC* insertion position, and individual barcode sequence (hereafter called TSS, *LUC*, Barcode).

### Histogram (A): read depth distribution (Figure S2a)

We firstly made a cumulative histogram by read-tags (TSS, *LUC*, and Barcode) for each experimental replicates. The histogram showed that 50 - 60% of tags appeared only one time in the results, which indicated that a significant amount of tags were erroneously generated due to the errors on either variable; TSS, *LUC*, or Barcode.

### Histogram (B): for Barcode validation (Figure S2b)

In order to elucidate erroneous barcode sequences from the results, we calculated the read occupancy of each Barcode in the associated locus (TSS and *LUC*). The histogram showed that low frequent Barcode species occupied 50 - 80 % of the reads that were mapped onto the same TSS-to-*LUC* loci.

### Histogram (C): for Barcode and TSS validation (Figure S2c)

Each *LUC* insert can have multiple TSSs. Hence, for each *LUC* insert, the genomic position of associated TSSs can be multiple, but their barcode sequence should be the same. In order to validate TSSs for individual *LUC* inserts, we calculated the read occupancy of each Barcode species in the associated *LUC* locus with ignoring their TSS locus. The histogram showed that among the reads that were mapped onto the same *LUC* locus, lower frequent Barcode occupied 60 - 75 % of them.

### Histogram (D): for *LUC* validation (Figure S2d)

Erroneous sequencing data can cause the miss-identification of the *LUC*-genome junction during the read processing step, which will shift the mapping result of the *LUC* locus with several nucleotides. Thus, in order to validate *LUC* insertion sites, we calculated the read occupancy of each *LUC* in the associated TSS with Barcode. The histogram showed that 10 - 20% of reads were mapped on different *LUC* locus among respective TSS-barcode sets.

Based on the above histograms, we collected tags when they fulfill following cases; (1) at least two counts (based on the histogram (A)), and (2) reads of which occupancies in histograms (B), (C), and (D) are more than 70%. If any above conditions were not satisfied, those reads were discarded regarded as erroneous ones.

---

**Figure S1.**
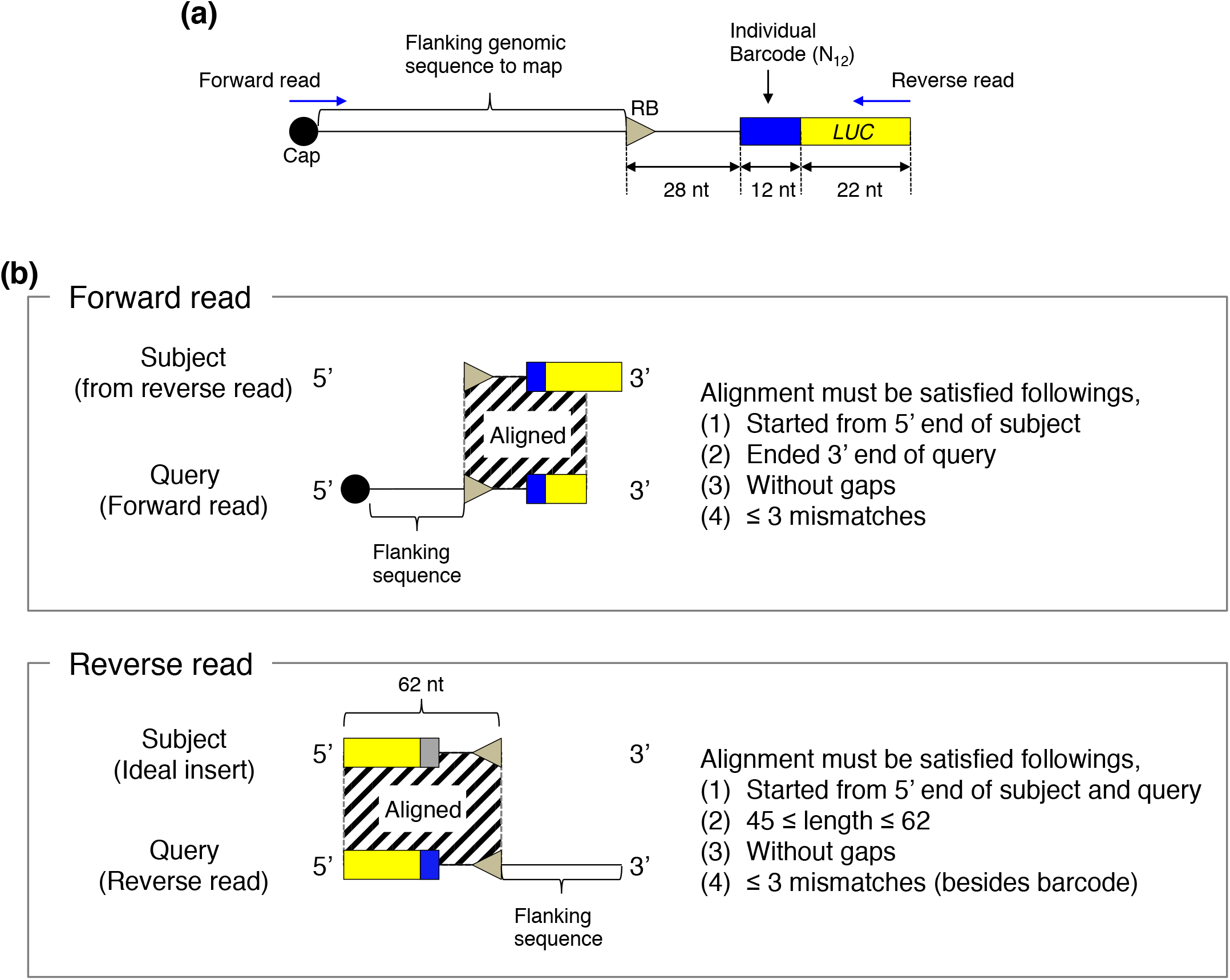
Schematic illustration of paired-end read processing before mapping. (**a**) Sequencing library design. (**b**) Detection scheme of *LUC*-genome junction from forward (Top panel) and reverse (Bottom panel) sequencing reads by Nucleotide BLAST. Shaded areas indicate an ideally aligned region. For detail, see Methods S1.

**Figure S2.**
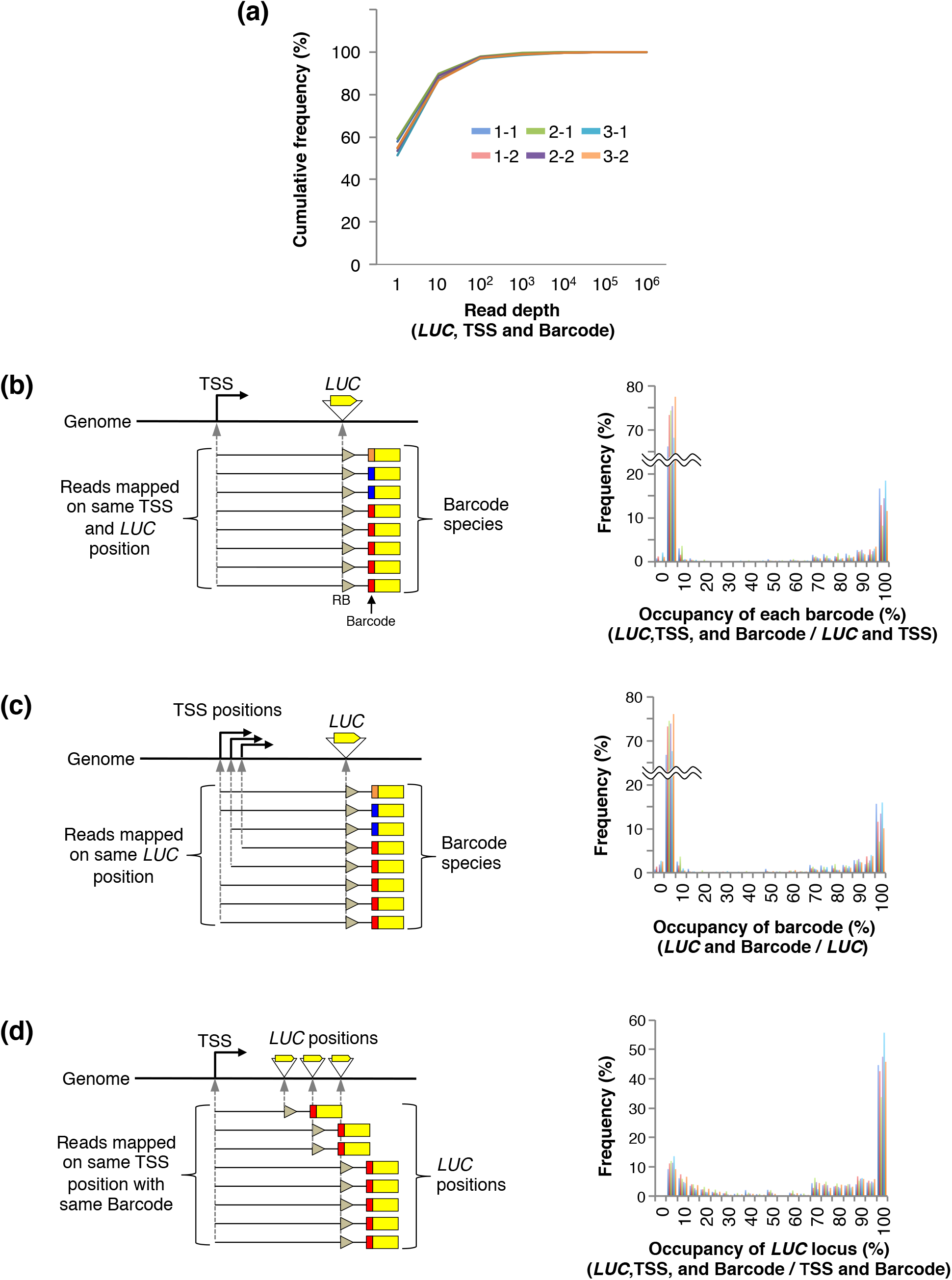
Mapping depth-based determination of genuine *LUC*-TSSs. (**a**) Cumulative frequency of *LUC*-TSS species (specified by TSS, *LUC*, and Barcode) according to their mapping depth in each experimental replicate. (**b**) Barcode validation. (Left) Barcode species and its abundances in the reads that mapped on the same TSS and *LUC* loci were calculated. Different color indicates different barcode sequences. (Right) Barplot shows the frequency of the occupancy of individual Barcode in the reads that mapped on the same TSS and *LUC* loci. (**c**) Barcode and TSS validation. (Left) Barcode species and their abundances in the reads that mapped on the same *LUC* locus were calculated. (Right) Barplot shows the frequency of the occupancy of individual Barcode in the reads that mapped on the same *LUC* loci. (**d**) *LUC* locus validation. (Left) Each frequency of *LUC* loci in the reads that mapped on the same TSS loci with the same Barcode sequences was calculated. (Right) Barplot shows the frequency of the occupancy of the individual *LUC* loci in the reads that mapped on the same TSS loci with the same barcode sequences.

**Figure S3.**
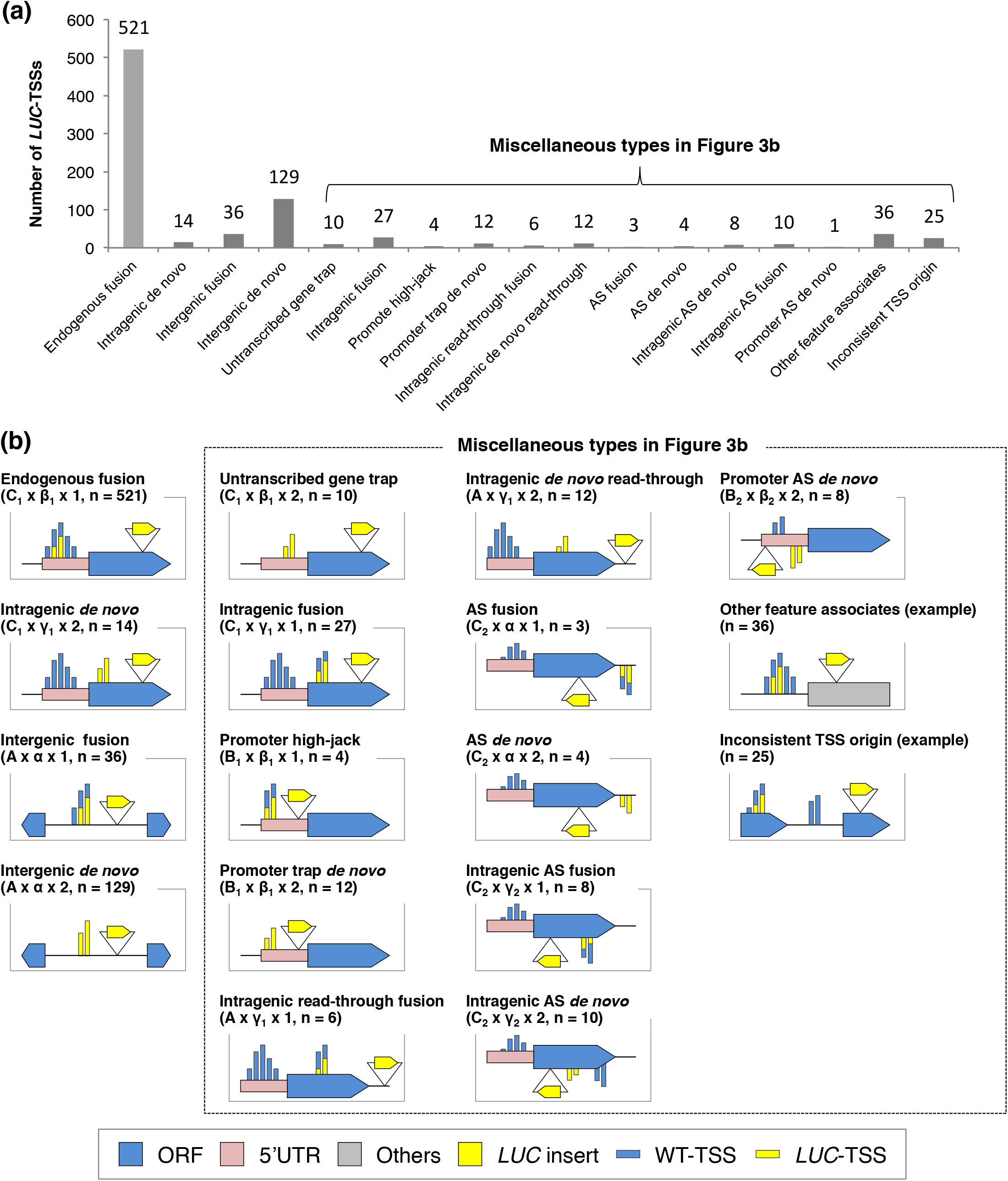
All determined *LUC*-TSS types. (**a**) The number of *LUC*-TSSs classified according to Figure 3a. AS: antisense to reference. (**b**) Schematic illustration of each type of *LUC*-TSSs. ‘Other feature associates’ and ‘Inconsistent TSS origin’ types represent an example of each type.

**Figure S4.**
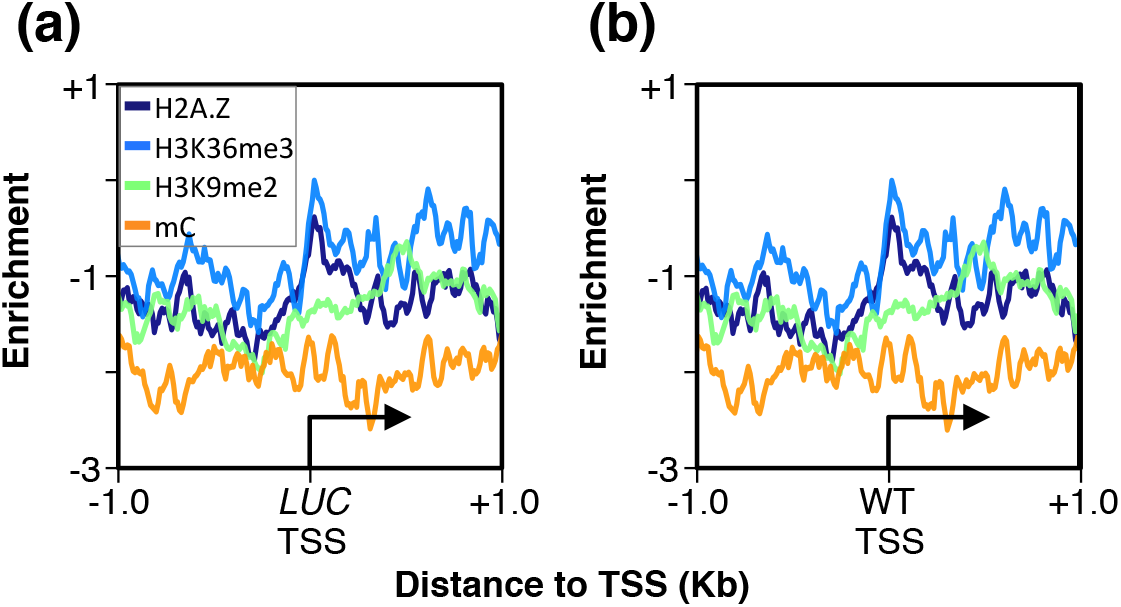
Distribution of epigenetic marks around the TSS of Intragenic *de novo* type. (**a and b**) Distribution profiles of H2A.Z, H3K36me3, H3K9me2, and methylated cytosine (mC) in WT cells, within +/- 1.0 kb of the TSSs of (**a**) Intragenic *de novo* type (from Figure 4c), and (**b**) corresponding trapped endogenous genes.

**Figure S5.**
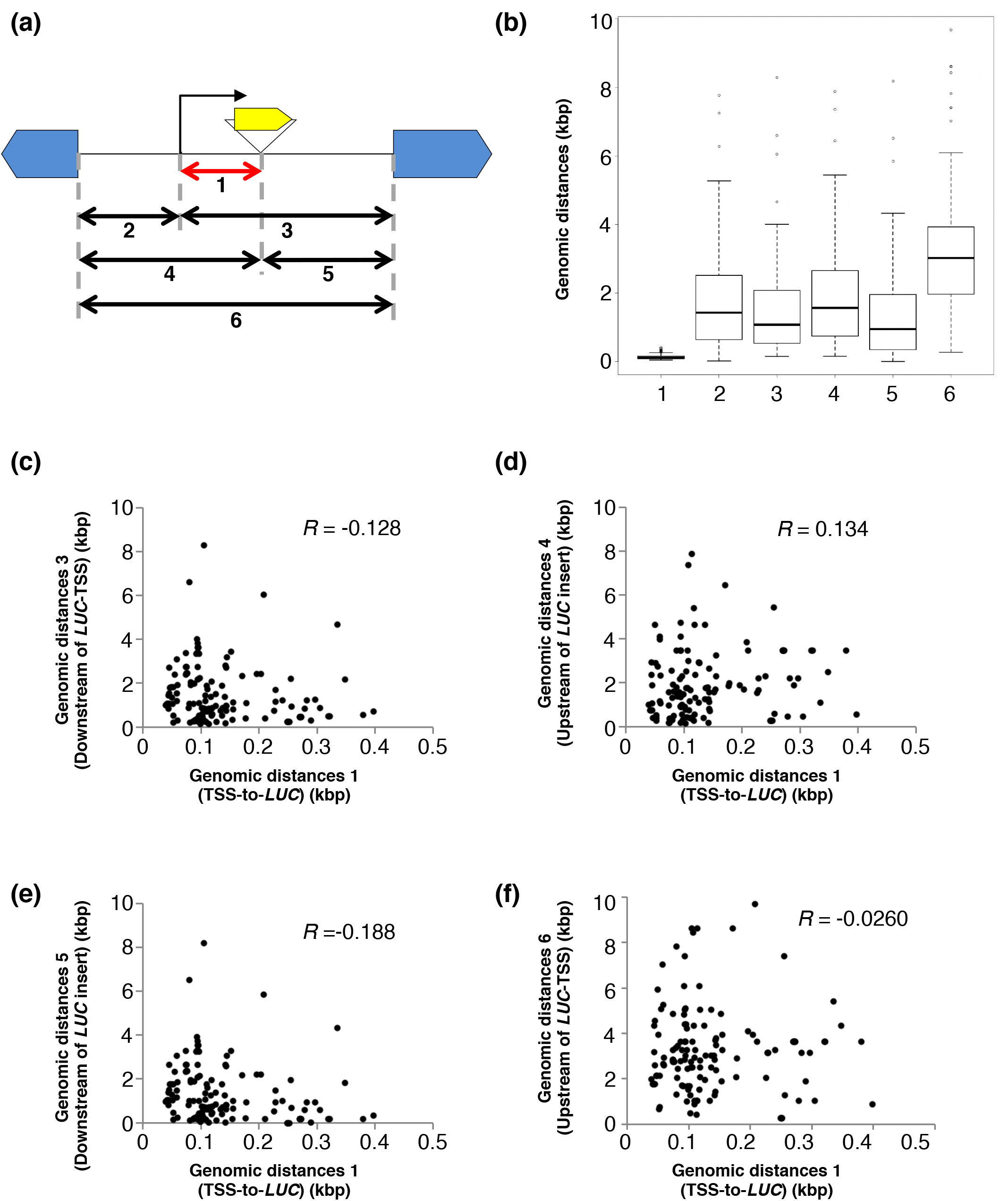
Comparison between the length of *de novo* TSS region and that of the corresponding intergenic region. (**a**) Schematic illustration of distances to be compared; 1: *TSS-to-LUC*, 2: from upstream neighboring gene to TSS, 3: from TSS to downstream neighboring gene, 4: from upstream neighboring gene to *LUC* insertion site, 5: from *LUC* insertion site to downstream genomic feature, and 6: width of the intergenic region. (**b**) Boxplots show the distribution of each distance classified in (a). (**c–f**) Scatterplots show a comparison between genomic distances of TSS-to-*LUC* (Intergenic *de novo* type) and region 3 (c), 4 (d), 5 (e), and 6 (f), of which regions were classified in (a), respectively. *R*, Pearson’s product-moment correlation test.

**Figure S6.**
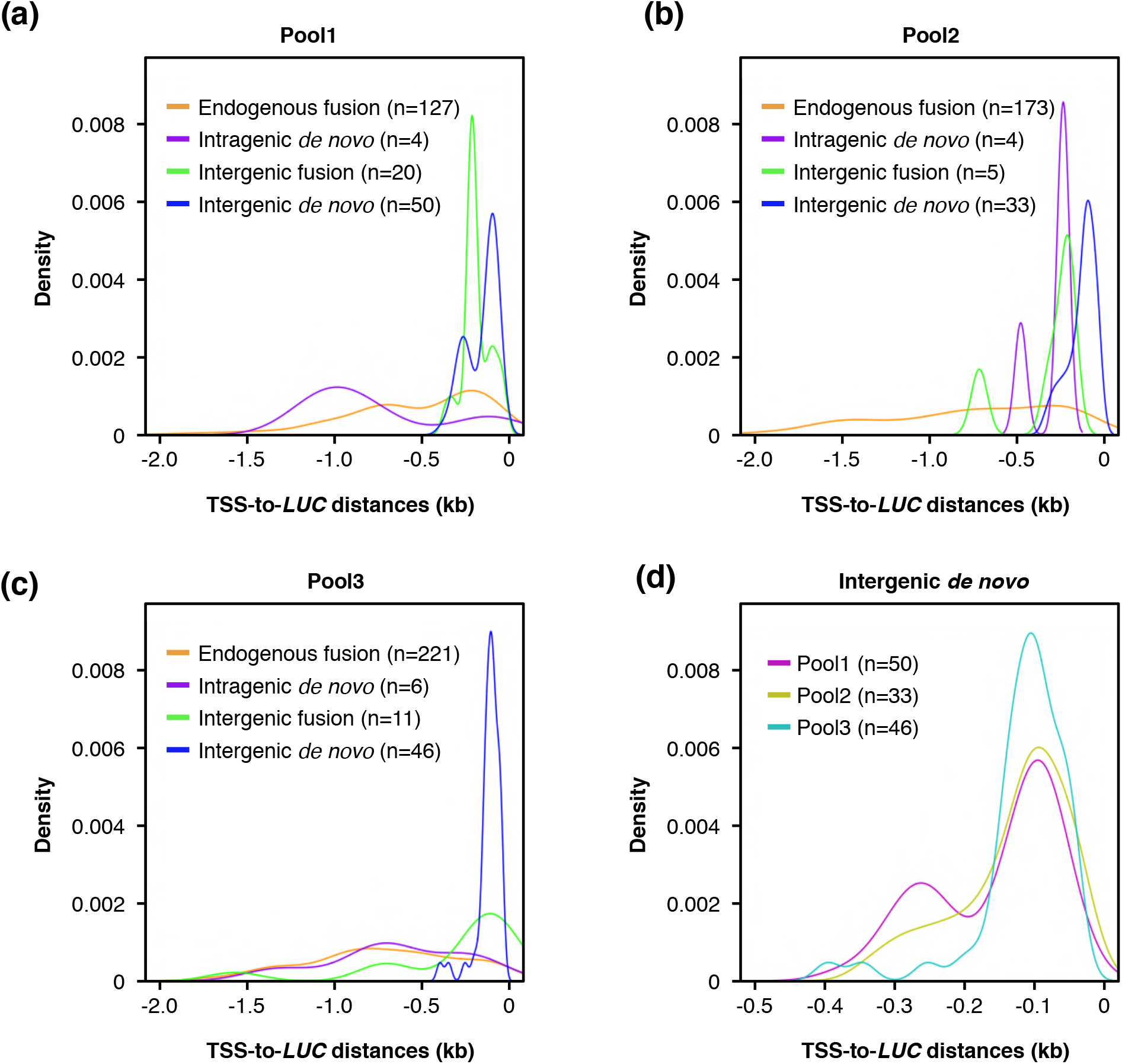
Distribution profiles of TSS-to-*LUC* distances in biological replicates. (**a–c**) Density plot shows distribution of distance between TSS and *LUC* insertion site *(TSS-to-LUC* distance) in each *LUC*-TSS type in distinct biological replicates; pool1 (a), pool2 (b), and pool3 (c). (**d**) Distribution profiles of TSS-to-*LUC* distances of intergenic *de novo* types in each biological pool.

**Figure S7.**
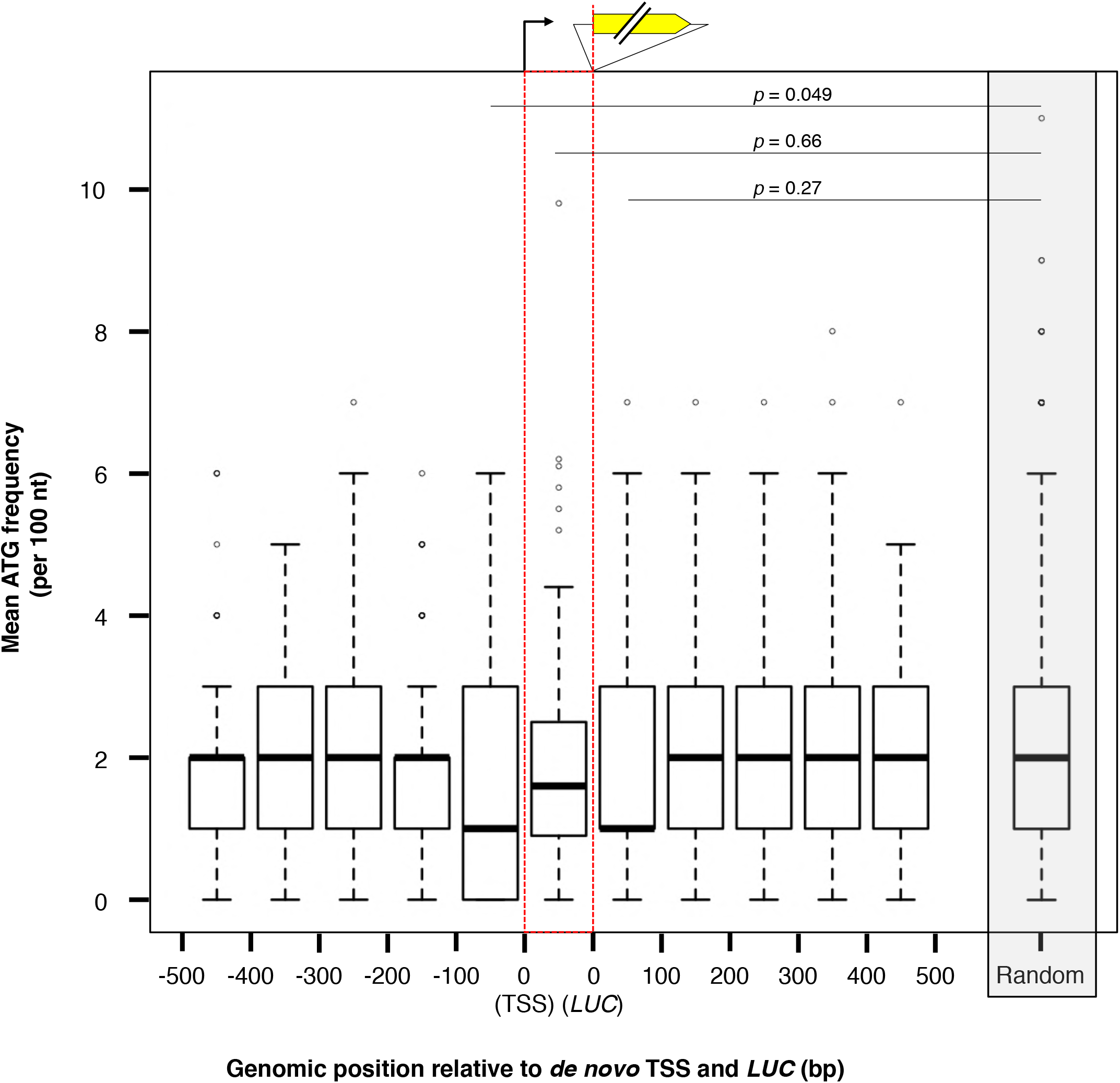
Initiation codon frequency around *de novo* TSS region. Boxplots show the mean frequency of initiation codon (ATG) around *de novo* TSS regions in 100 bp windows. The ATG frequency at *de novo* TSS region was normalized to per 100 bp, and was indicated by red dotted box. Grey shaded box indicates the mean ATG frequency of the 10,000 times bootstrap sampled intergenic region. *P*-values of the Wilcoxon rank sum test are indicated on the plot.

**Figure S8.**
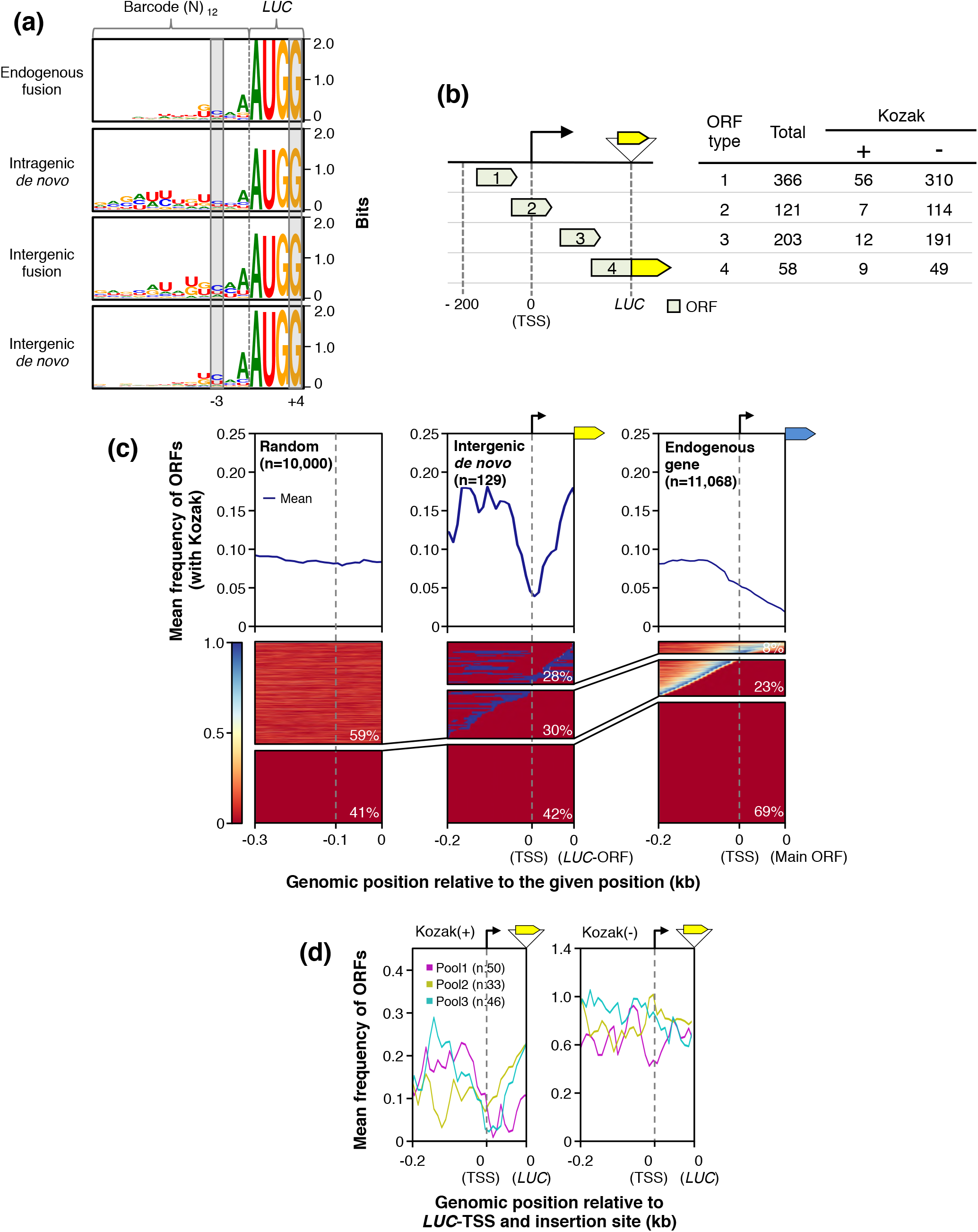
Distribution profiles of Kozak sequence around *LUC* insertion loci. (**a**) Sequence logos of barcode sequences of *LUC* inserts classified according to Figure 3a. Positions of Kozak motifs are indicated by grey boxes. (**b**) Putative ORFs were classified into four types; (1) – 200 bp upstream from *de novo* TSS, (2) over the de novo TSS, (3) between *de novo* TSS to *LUC*-ORF, and (4) fusion with *LUC* ORF. The right table shows the number of putative ORFs found. (**c**) Metaplot and heatmap of distribution profiles of with Kozak-motif within 0.3 kbp of randomly sampled intergenic regions (left panels), the region from 0.2 kbp upstream of the Intergenic *de novo* TSS to its *LUC*-ORF (middle panels), and the region from 0.2 kbp upstream of the TSS of endogenous protein-coding gene to its main ORF (right panels). As for the regions between individual *de novo* TSS and *LUC*-ORF, their real lengths were varied according to individual sites. Therefore, their individual lengths were normalized to 100 bp when calculating ATG frequency. *Arabidopsis* genes with introns in the 5’-UTR were excluded from the analysis. Gray dotted lines indicate TSS positions. (**d**) Metaplot of distribution profiles of pre-existing ORFs with (left panel) or without (right panel) Kozak sequences. *De novo* TSSs in the different biological pools were analyzed separately as in (c).

**Table S1.**
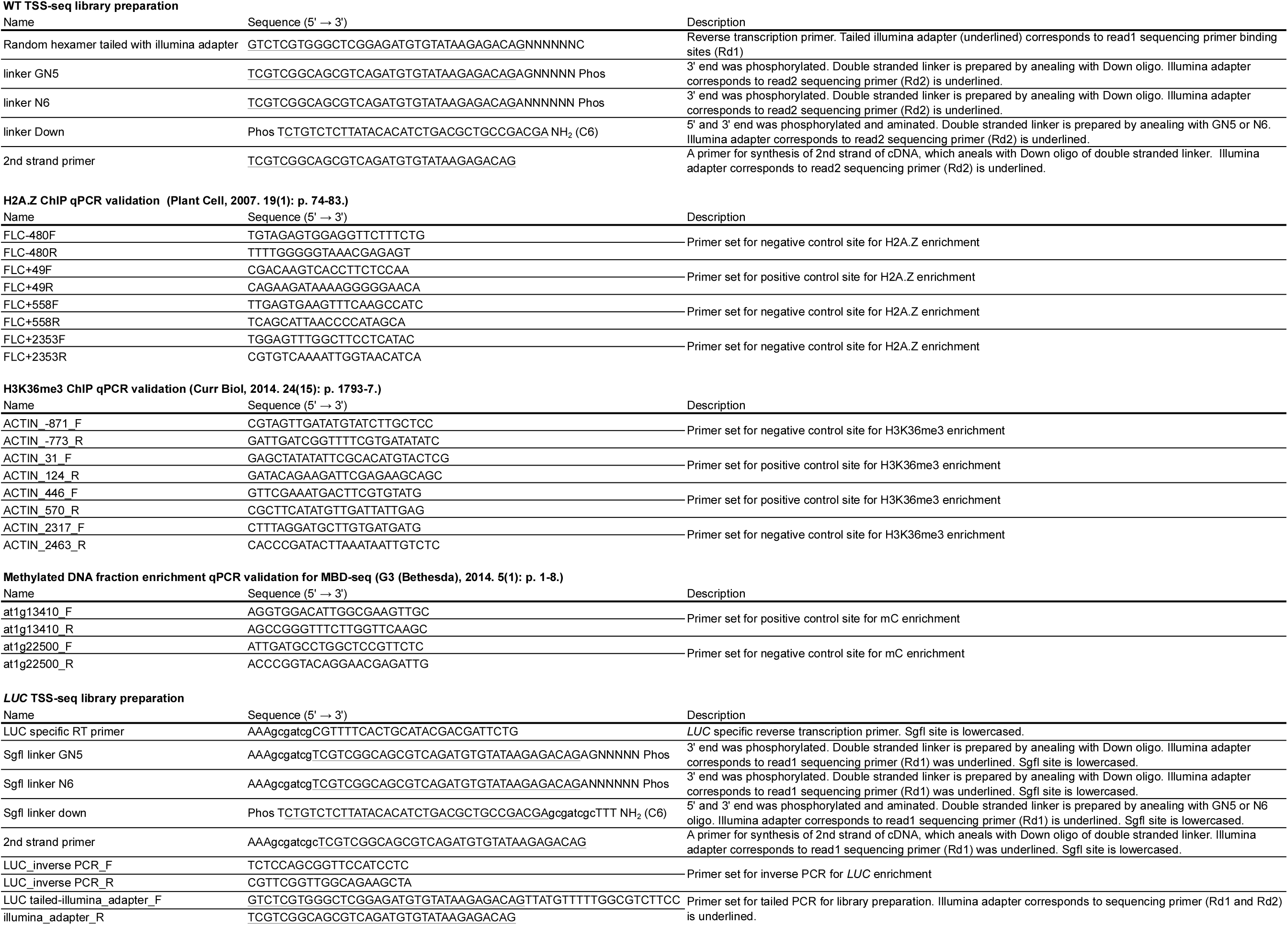
Information of All DNA oligos used in this study.

